# Mora-ERP-based RNN-Transformer for decoding single-trial EEGs during silent Japanese speeches

**DOI:** 10.1101/2024.11.18.624218

**Authors:** Toshimasa Yamazaki, Saori Kudo, Sei-Ichiro Kamata, Satoshi Fujii, Sho Tsukiyama, Tetsushi Yata, Shunsuke Aoki

## Abstract

We developed a method for decoding single-trial electroencephalography (EEG) during silent Japanese speeches. In order to cope with problems that there would be always noises in single-trial EEGs, a recurrent neural network (RNN) was used which could reproduce signals under noises. Each of silent-mora-related potentials and the single-trial EEG minus the event-related potential (ERP) were assigned to the signal and the noise, respectively, with reference to the averaging principle. Next, in our Transformer, dot product between the RNN output and the single-trial EEG after positional encoding then Softmax with Loss yielded probabilities of moras, each of which consists of silent Japanese words, phrases or part of sentences. The present decoding was completed by tracing the maximal probability at each block representing time. Average mora error rates (MERs) on pretrained and validated performances for the patient was as low as 1.5 % and 0 %, respectively. The performance for the testing would be refined by many single-trial EEGs during silent Japanese speeches obtained by EEG Web interfaces. This method might be applied to other “mora” languages.

---

In the last three decades, speech has been decoded from human brain signals invasively^1-7^ using ECoG (electrocorticography) and non-invasively. The latter utilizing such as EEG (electroencephalography)^8^, MEG (magnetoencephalography)^9^ and fMRI (functional magnetic resonance imaging)^10-12^ have been attempted. Especially, the EEGs had been decoded during silent English words^13^, isolated phoneme^14,15^, monosyllable^16^, and silent Japanese 2-mora (mentioned later) words^17^. Recently, the Transformers^18^ have been applied to MI-BCI^19^ and EEG-to-text decoding^20,21^, the latter accuracy does not remain so good.

Such EEG-based decoding should be more widely adopted, because it does not require invasive neurosurgery but also is compact, portable, economic and suitable for quality of life (QOL)^22^. However, decoding of single-trial EEGs have not been so much in advance due to the low signal to noise ratio of raw EEG data (for example, see ref.^23^). Averaging, bringing event-related potentials (ERPs), is one of solutions for the problem, but not suitable for real-time decoding. The principle of averaging is based on that any single-trial EEG is assumed to be signal plus noise whose probabilistic distribution has the mean of zero. Arecurrent neural network (RNN) has been known to reproduce spatiotemporal patterns to be memorized under the noise^24^. Given a single-trial EEG with any task, if the noise could be assigned to the EEG minus the signal, that is, the ERP, according to the averaging principle, the RNN output might yield an estimate of the average task-related potential under the noise.

Japanese has syllable-timed rhythm and is called a “mora” language^25^, and the mora is the phonological unit of word production^26,27^. Moras, constructing any word or phrase, are positionally governed by the sonority sequencing principle^28^, and the mora is discriminable according to the degree of the sonority. Therefore, all the words or phrases in Japanese can be represented by the moras. In this study, the inputs to the above RNN are each of average ERPs during silent Japanese moras as signals, and single-trial EEGs during silent words or phrases minus the signals as the noises.

The present neural network model has multi-head attention (MHA) structure in the Transformer^18^, where *Q* (query) is assigned to decomposed single-trial EEG, *K* (key) is obtained by the above RNN output for each silent mora ERP, and *V* (value) is given so that the *V* and Affine layer form LSTM (long short-term memory)^29^. The dot-product between the *Q* and the *K* indicates how much the *Q* included each *K*. The number of the heads is equal to that of the blocks. Next, “Softmax with Loss” layer, which calculates Softmax function and cross entropy error, is involved. Other loss functions can be applied such as MSE Loss and CTC (connectionist temporal classifier) Loss, where the latter Loss can train the neural network to output a sequence of symbols (in this case, moras and blanks) given unlabeled time series input^30^. Lastly, mora probabilities were generated as functions of block number. Tracing mora with the maximal probability at each block yields the decoding of single-trial EEG during silent Japanese speech.

## Results

Participants except for one female patient silently spoke moras (see Table 1^31^), words or phrases presented hiragana or katakana on the monitor, while the patient covertly spoke only words and phrases.

**Table 1.**
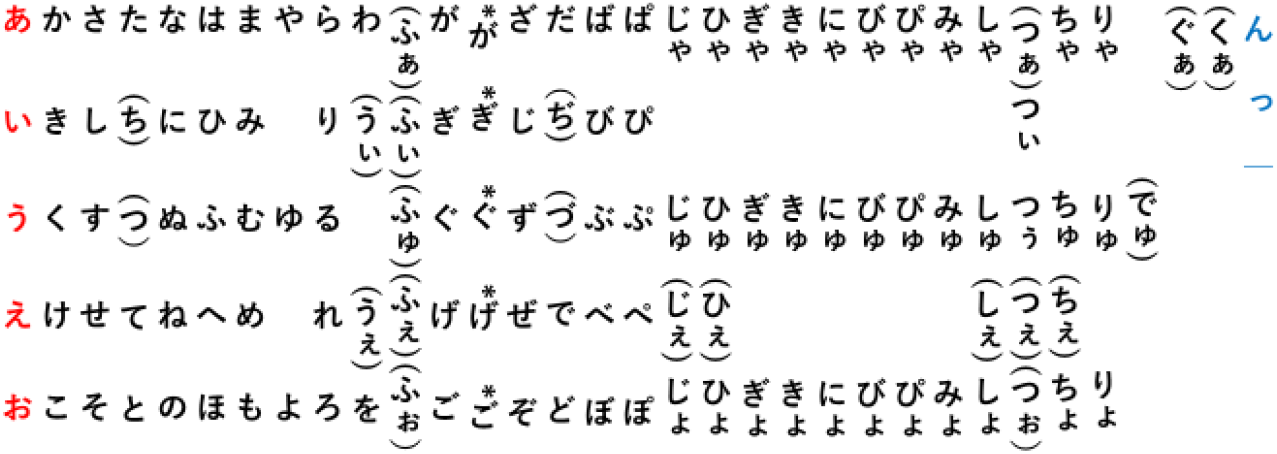
Japanese moras. The first column (red) corresponds to Japanese vowels, and three moras in the last column (blue), 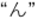, “っ” and “−” to a syllabic nasal, a double consonant and a major scale, respectively, the last two of which cannot be solely pronounced. Moras in parentheses are used only for foreign languages and interjections. *: nasal sounds of 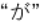 column

### Decoding pipeline

We begin with a brief description of the decoding pipeline, illustrated in Fig. 1, and described for details in the Methods. Single-trial EEG (electroencephalography) data were recorded from active electrodes or a Muse EEG headband^22^, while the patient silently spoke words, phrases or sentences. The data only at F7 or AF7 passed as input ones to an artificial neural network, which processes the data in three stages:

**Fig. 1.**
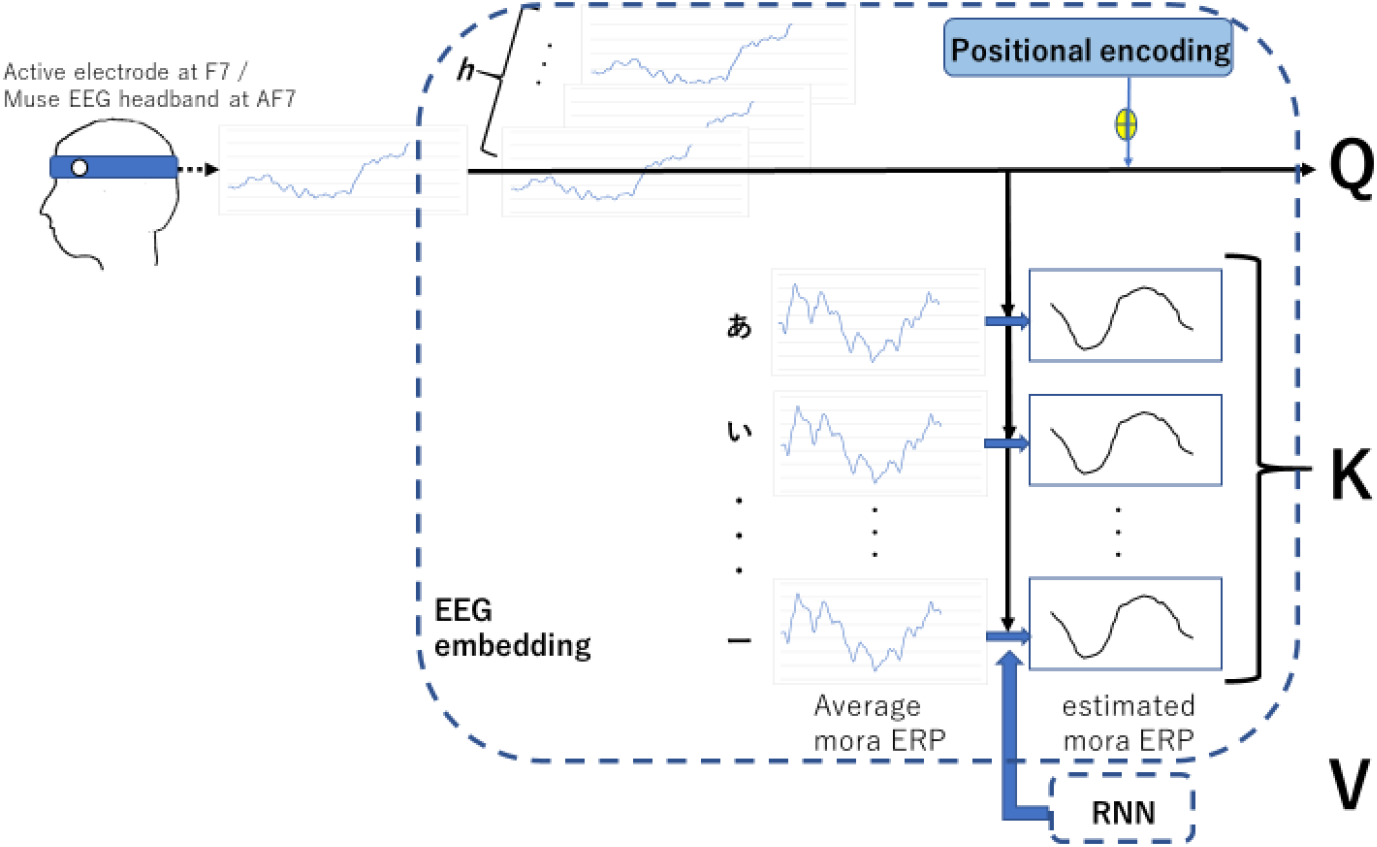
The decoding pipeline, a, EEG embedding and b, RNN.

**Fig. 1a.**
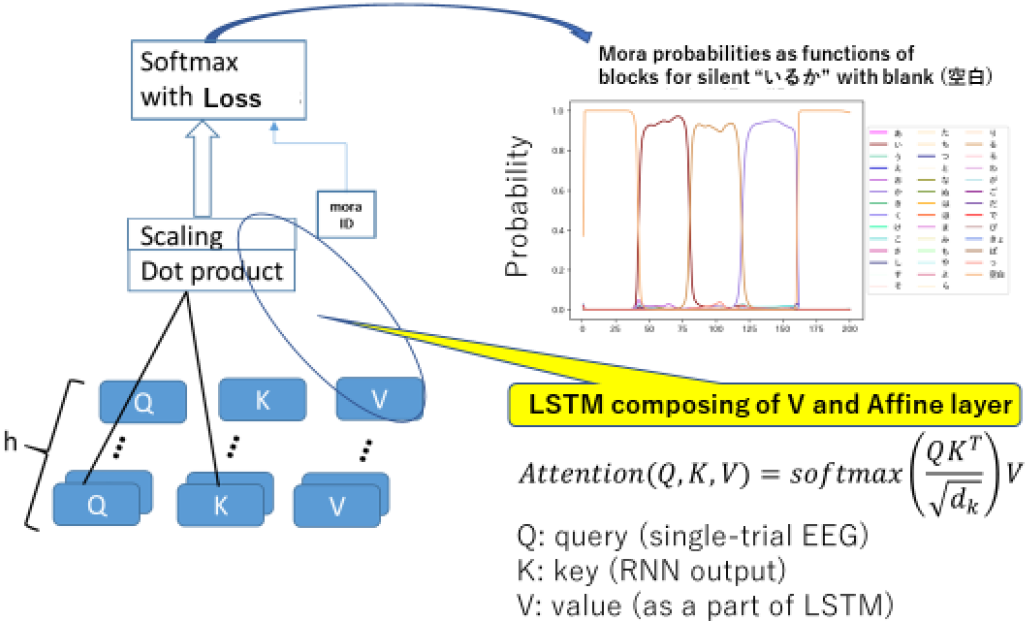
The decoding pipeline, c, Transformer.

1. **EEG embedding:** the EEG embedding refers to segmentation between 0-300 ms, including N1 and P2, then time-overlappingly blocking of the data.
2. **RNN:** using each of the blocked single-trial EEGs and silent-mora ERPs, under the specified noise the ERPs were estimated by a recurrent neural network (RNN).
3. **Transformer:** *Multi-head attention (MHA)*. Our model has MHA structure, where Q (query) is assigned to each of blocked single-trial EEG, K (key) is obtained by the above RNN output for each silent mora ERP, and V (value) is given so that the V and Affine layer form LSTM (long short-term memory)^29^. The dot-product between the Q and the K indicates how much the Q included each K. The relationship among Q, K and V is given by

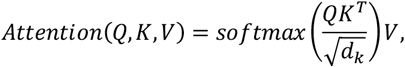

where *dk* is the dimension of Q and K. The number of the heads is equal to that of the blocks. Each block was blanching to the dot-product layer and the above RNN, as the self-attention in the Transformer.

*Softmax with Loss*. Next, “Softmax with Loss” layer calculates Softmax function and cross entropy error. Other loss function can be applied such as MSE Loss and CTC Loss, where the latter Loss can train the neural network to output a sequence of symbols (in this case, moras) given unlabeled time series input^30^. Lastly, mora probabilities were generated as functions of block number. Tracing mora with the maximum probability at each block yields the decoding of single-trial EEG during silent Japanese speech.

The present Transformer is trained to make it assign high probability to target moras at each block, using teaching signals. All training proceeds by stochastic gradient descent via backpropagation.

### The RNN performance

Fig. 2a shows the output of the present RNN (black curve), where the signal was assigned to an average ERP (grey curve) during silent Japanese vowel “あ” and the noise to one single-trial EEG during the silent “あ” minus the signal. Fig. 2b corresponds to the case where the signal was given by an average ERP during silent Japanese word “うたご*え” (/utagoe/) (meant by singing voice in English), where “ご*” indicates nasal sound of “ご”. These figures demonstrate that the present RNN led us to the reconstruction of the average ERPs as the signals^32^.

**Fig. 2.**
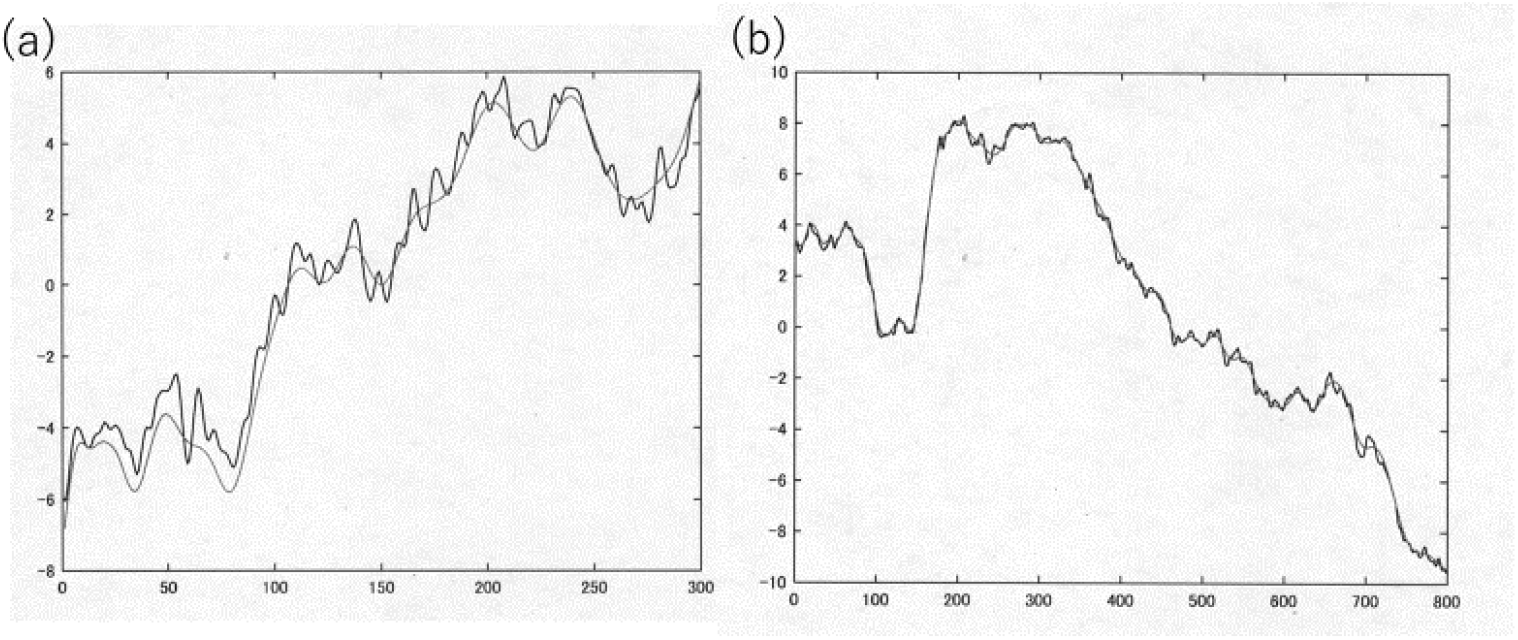
Predicted and averaged ERPs for a mora “あ” (A) and a word “うたご*え” (B).

### Decoding performance

Here and throughout, we quantify performance with the average (across all trained, validated and tested words, phrases and sentences) mora error rate (MER); that is, the minimum number of deletions, insertions and substitutions required to transform the predicted words or phrases into the true ones, normalized by the number (N’) of moras between two spaces at both ends. If the number (N) of moras consisting of words or phrases is less than N’, there would be insertions and substitutions, if N more than N’, deletions and substitutions. Thus, the MER for perfect decoding is 0 %, its example is that moras, which form words, phrases or sentences, appear in turn between the two spaces, as shown in Fig. 3. This figure exemplifies a decoding result of silent “がっ こう (school in English)”. The right side shows candidate moras including “blank (空白)”. Note that an ERP for “っ” is obtained by substracting “あ”-ERP from “あっ”-one, because of “が” involving a vowel “あ”.

**Fig. 3.**
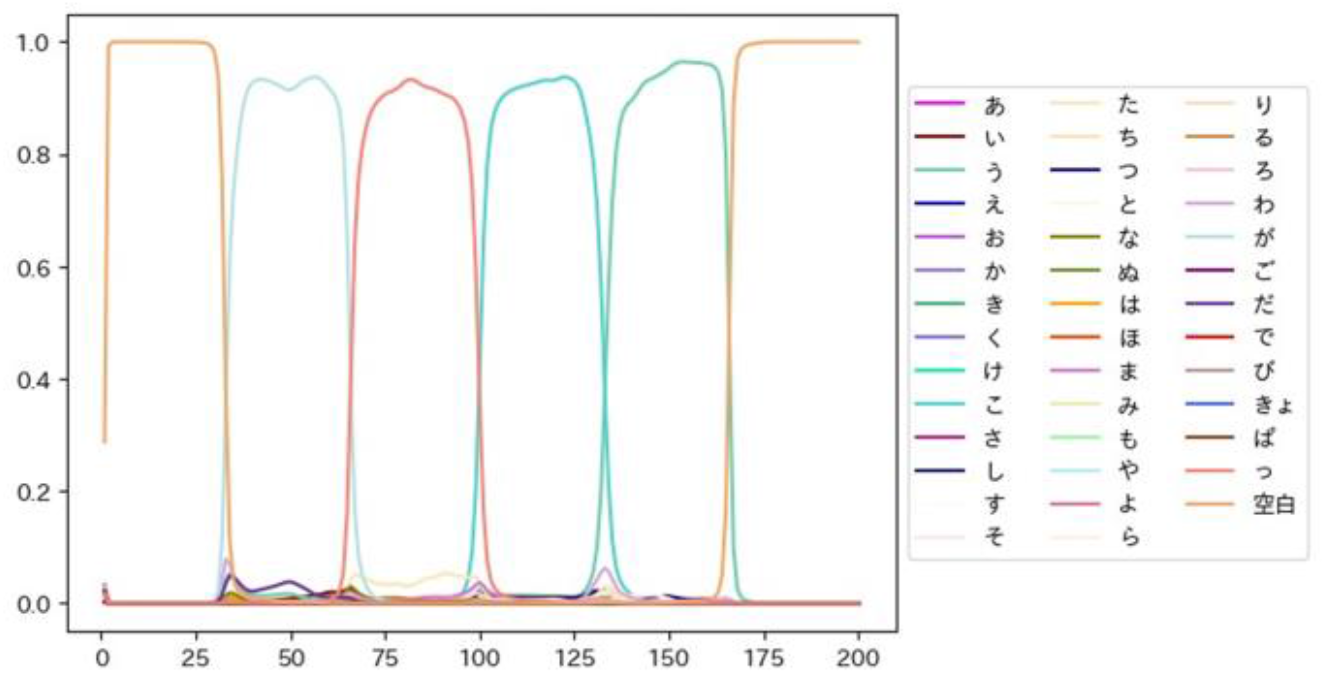
Decoding result of silent がっこう (school in English)” with blank (26).

Table 2 shows MERs for up to 41 trained (validated and tested) words, phrases and (part of) sentences by the patient. The mean MER for the patient as the pre-training performance is approximately 1.5 %. The performance at the validation phase, that is, the same words but other trials as the pre-trained, was that average MER was 0 for 8 silent words. However, the performance at the testing level was not good (MER = 50 %) due to silent Japanese speeches of the limited number of some dozens. Note that our RNN yielded better performance than silent-mora-related potentials.

**Table 2.**
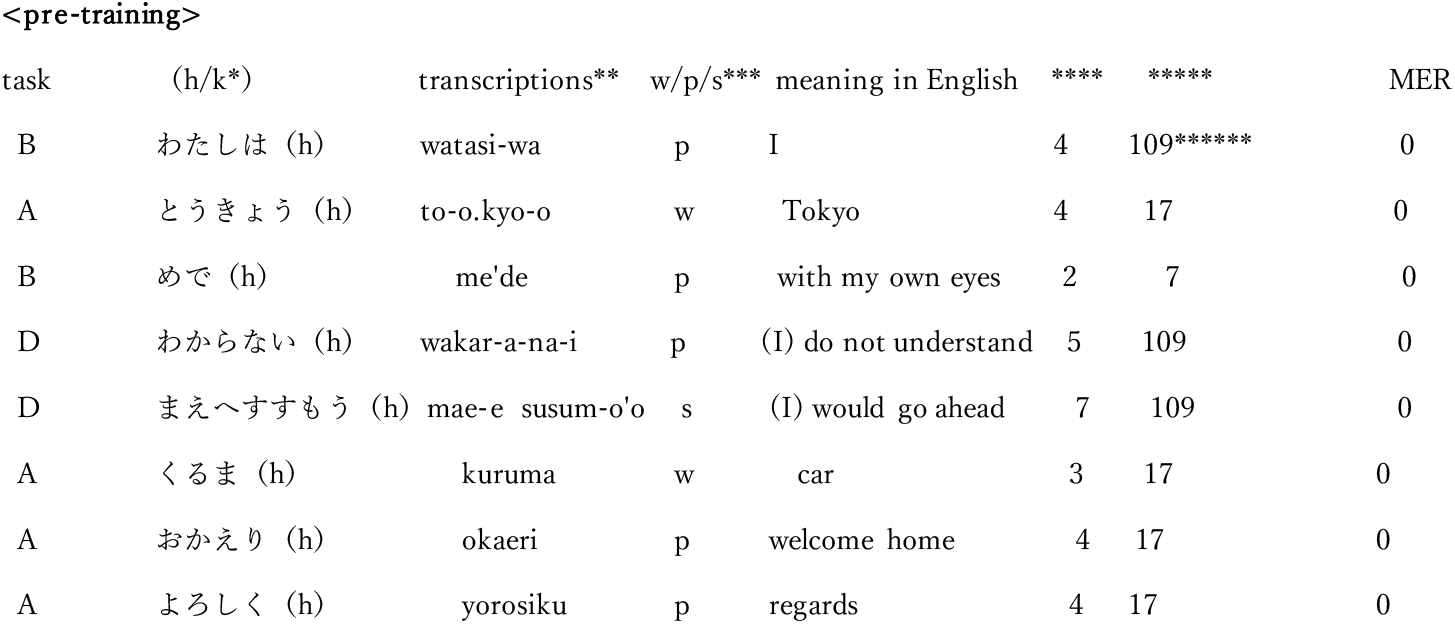
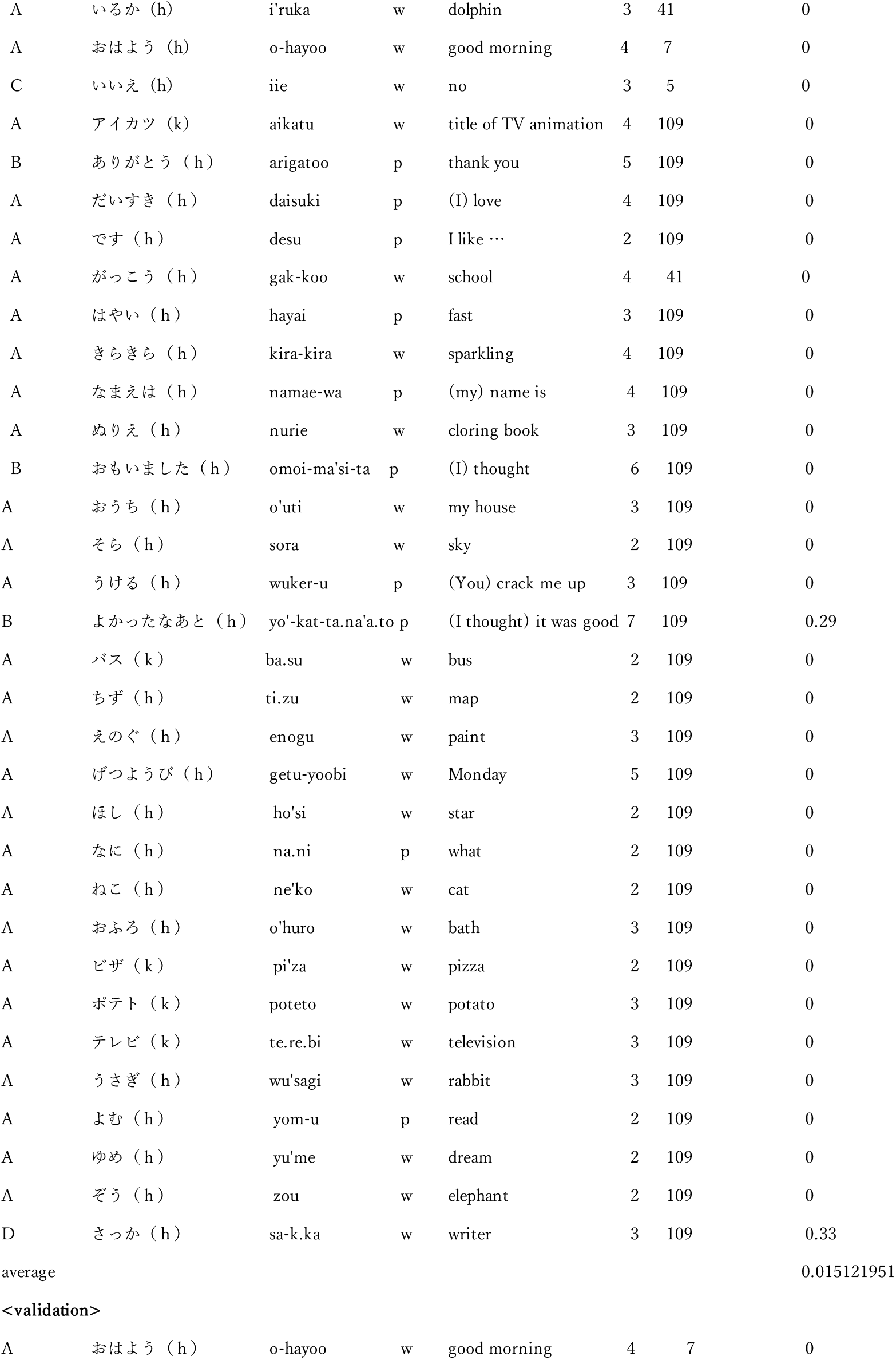
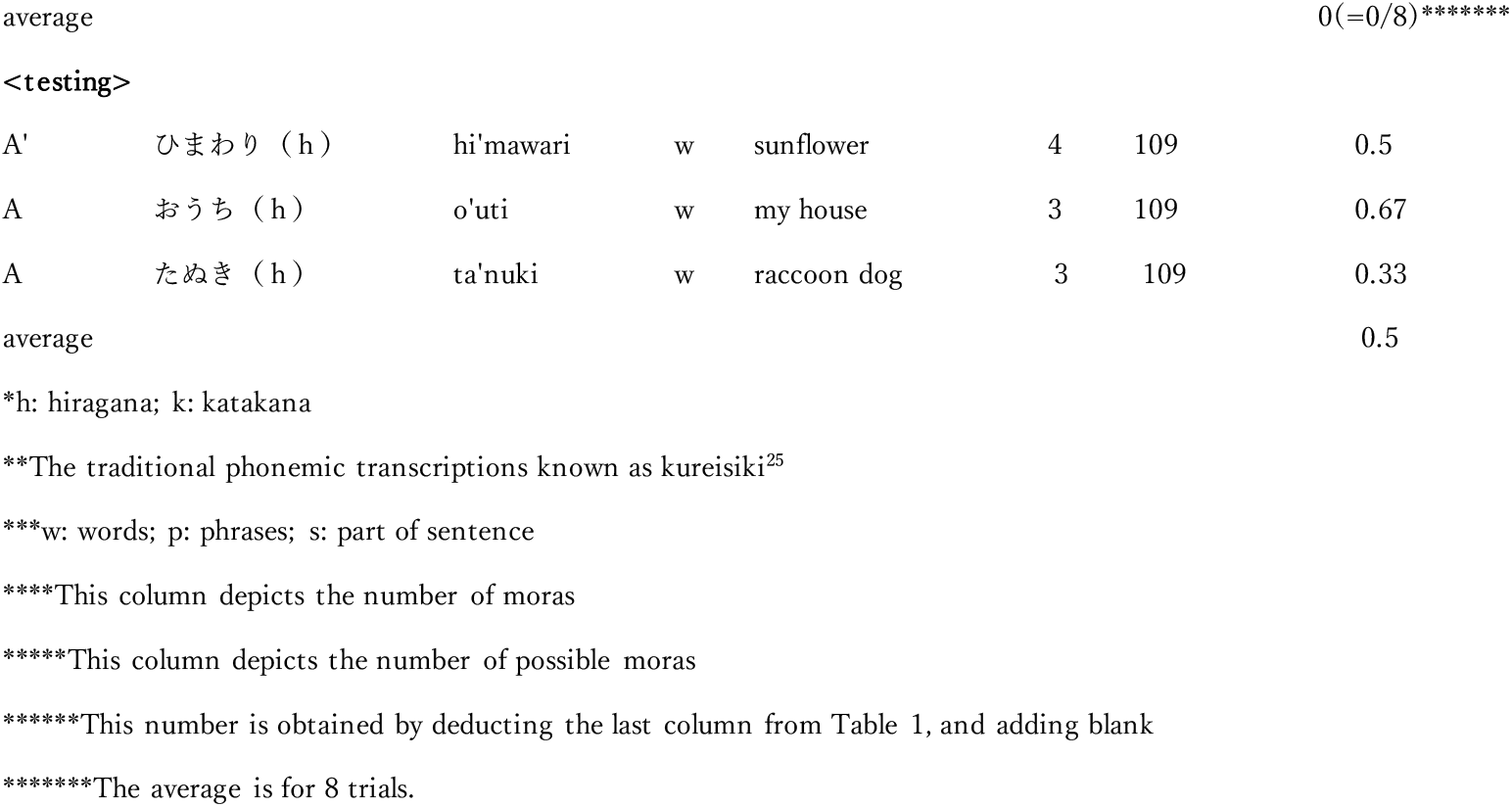
Performances for the pre-trained, validated and tested trials.

### Why N1 and P2 only at F7 were analyzed

During both the overt and covert production of phonemes, words and phrases auditorily^33-36^ and visually^2,27,37-39^ presented, N1 and P2 similarly appear in scalp-recorded EEGs. In our preliminary experiment, silent speech trials of both 800 pseudowords and each of 800 moras producing the pseudowords which are any pair of moras without meanings yield similar N1 and P2 (see Fig. 4^40^), as shown in Fig. 4.

**Fig. 4.**
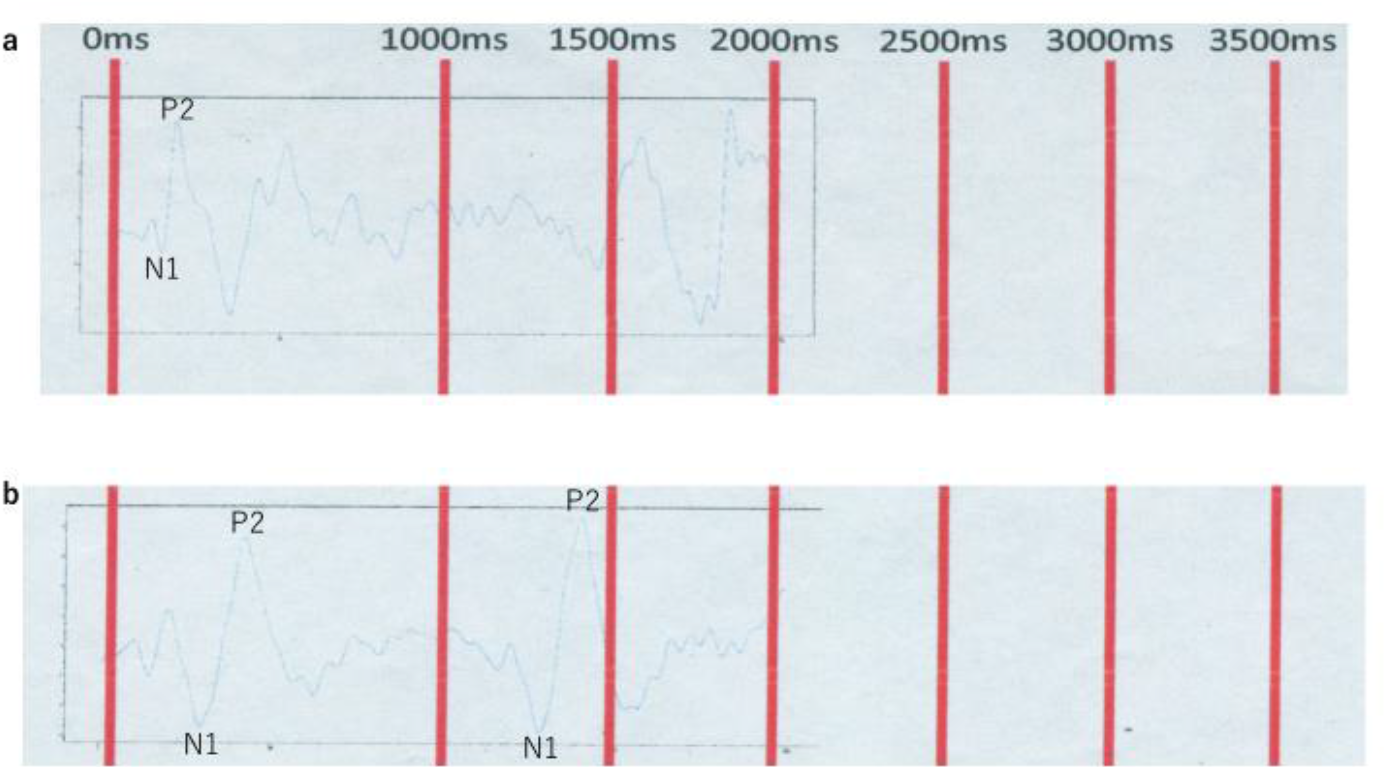
Silent-speech-related potentials, **a**. ERP during silent word whose moras were simultaneously presented, **b**, ERP during each of silent moras delayed by 1 second, where the moras produce the word.

P2 might reflect a ‘temporal-matching response’ in terms of whether the audible phoneme matched its inner phoneme, which would yield efference copies (corollary discharge) ^34,36^. The efference copies are the crucial concept driving internal forward model with feedback^41^, involving self-monitoring^42^ and as state feedback control^43^. For 19-ch ERPs, z-scored ERPs at F7, C5 and T7 yielded one cluster with unsupervised hierarchical clustering (Fig. 5^39^), which might indicate the flow of the efference copies. And this component including P3 might reflect the phonological processing in word production^2^. Moreover, green vertical dotted lines in this figure indicate speech onset estimated by EMGs (electromyograms) when actually speaking. This finding might show that N1 is not directly related with the phonological processing.

**Fig. 5.**
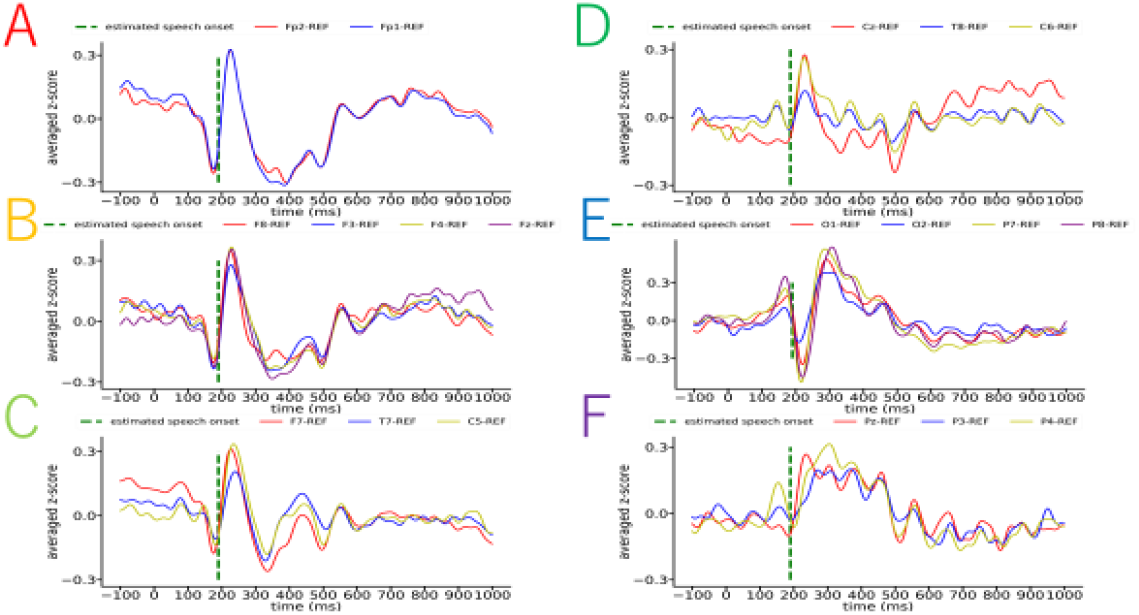
Unsupervised hierarchical clustering for 19-ch z-scored ERPs

The N1 is included in ELAN correlated with rapidly detectable word-category errors^44^, and might reflect a lexical access from a graphemic code in word reading^2,45^. Moreover, unilateral (left) and bilateral N1s reflect attention-related and more automatic orthographic process, respectively^46,47^.

The previous neurocognitive studies have proposed that the left inferior frontal gyrus might lie at the phonological end of nonlexical grapheme-phoneme (or hiragana or katakana-mora in this study) conversion processes in reading^48^, and that the left posterior superior temporal lobe might be involved in lexical phonological code retrieval^42^. Fig. 6^49,50^ shows average ERPs during silent “はい” (yes meant in English) and “いいえ” (no in English) at F7, where the number of summations was 1000. From this figure, amplitude differences in P1 and P2 between “はい” and “いいえ” might lead to those in vision (graphic) and phonology, respectively, while N1s are the same. Thus, these findings yielded the reason why the interval to be analyzed was 0 to 400 ms only at F7 in this study.

**Fig. 6.**
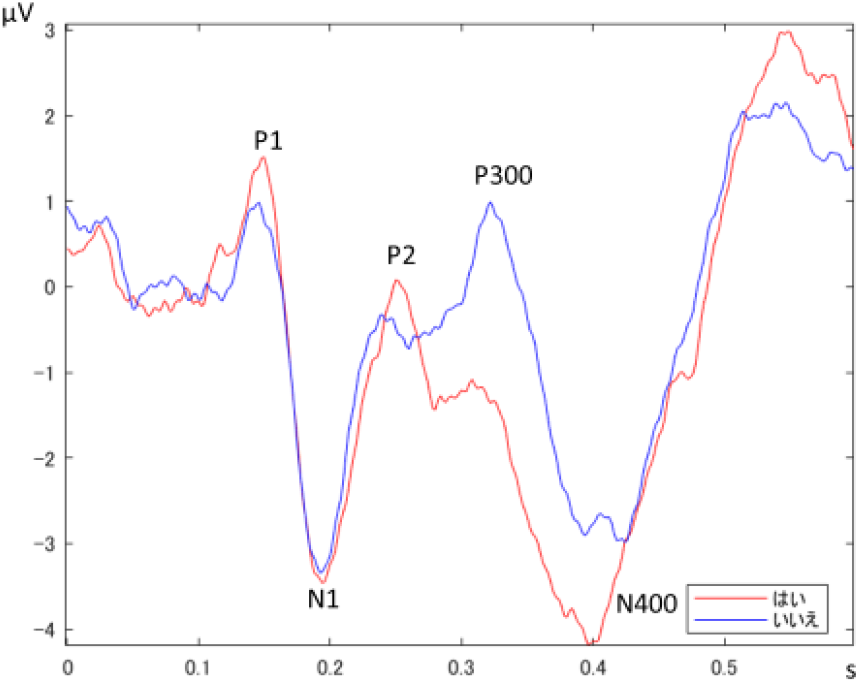
Silent “はい“ and “ いいえ”related potentials, where the number of summations is 1,000.

### Hyperparameters

We made a list of all hyperparameters in our neural network model, as shown in Table 3. Each value yielded the best performance so far. The summation number of silent mora ERPs might be crucial for the performance, while the number of possible moras not. Moreover, hyperparameters concerning blocks should be strictly investigated, because of sampling frequencies of EEGs.

**Table 3.** Hyperparameters in this model.

| Architecture | Hyperparameter | values** |
| --- | --- | --- |
| EEG embedding | block size | 400 ms |
|  | distance between blocks | 4 ms*** |
|  | number of blocks | 100 |
| RNN | number of averaging for silent-mora-related potentials | 100 |
| | $\tau$ (unit time constant) | 0.001 |
| Transformer | number of possible moras | 109 |
|  | Loss*/MSE Loss/CTC Loss | MSE Loss |

\*In general, Loss function is described by mutual information
\*\*Each value yielded the best performance
\*\*\*The distances of 4 ms and 1 ms correspond to EEG sampling frequencies of 256 Hz and 1 kHz,
respectively

## Discussion

We have shown that silent speech in Japanese can be decoded from single-trial EEGs at the level of words, phrases and part of sentences, with MERs as low as 1.5 % at the pre-training level.

One of the most serious problems for single-trial-EEG-based BCIs is that the low S/N ratio of raw EEG data. We used an RNN which can reproduce patterns (signals) to be memorized under noises, where here averaged ERPs during silent moras and the single-trial EEG minus each of the ERPs were assigned to the signal and noise, respectively. Our RNN utilized the single-trial EEG minus the ERP instead of Gaussian noises. Although this use may be supported at “The RNN performance”, more strict examination will be needed.

The present study paid attention to that Japanese is a mora language. Therefore, this method might be able to applied to other mora languages such ancient Greek and Hindi, where there would be many moras which can be solely spoken.

Finally, we should construct EEG Web interfaces^51^ with a MUSE EEG headband, where many single-trial EEG data from even patients without fatigue can be free to be measured and analyzed.

## Methods

Six healthy, right-handed student volunteers (one female) between the ages of 22 and 23, and one female patient aged 15 with spinal muscular atrophy (SMA) type I participated in this experiment, where the patient from 7 years ago. This study was conducted at Kyushu Institute of Technology (Iizuka, Japan), Information, Production and Systems Research Center, Waseda University (Kitakyushu, Japan) and the patient’s house, and approved by the Ethics Committee for Human Subject Research, Faculty of Computer Science and Systems Engineering, Kyushu Institute of Technology, and by the Academic Research Ethical Review Committee, Waseda University.

### Task

The students mainly made covert product of moras, while the patient silently spoke words, phrases and (part of) sentences according to the following tasks. The tasks consist of the following four (A, B, C, D), where the procedures are the same except for Tasks B and C.

*Task A*. First, a fixation point marked “+” was presented for 2 s on a computer monitor which is located 77 cm away in front of the students, while the monitor was located diagonally upward 77 cm away from the patient. Next, when a Japanese mora, word, phrase or part of sentence in hiragana or katakana to covertly speak was presented for 2 s, the participants silently speak the mora, word, phrase or part of sentence as soon as possible. In case of the silent mora, only one student participated. The number for averaging was 30 to 100 so that N1 and P2 clearly appear in the ERP. The number of moras consisting of words, phrases and sentences used in this experiment ranges 2 to 7.

*Task B. (Diary-reading tasks)*. The patient was instructed to silently read every phrase in test excerpts from her diary, written in papers. The papers are held upwardly inclined about 50 cm away from her by the experimenter. The diary had been written by her with special computer keyboard controlled by her line of sight.

*Task C. (Question-and-answer task 1*^49,50^). EEG measurement starts with a click of highly sensitive mouse by the patient. Then, one question in Japanese is displayed on the monitor for 5 s. The monitor is located diagonally upward 70 cm away from her. Silent answers of yes or no are continuously iterated five times. Each answer needs two seconds. Twenty questions are prepared so that true yes:no is 1:1. For example, in case of “do you come from Fukuoka prefecture?”, the answer is “yes”.

An average ERP for each answer is decomposed into temporally overlapping blocks, and principal component analysis (PCA) is applied to these blocks. The first and second principal components are features. The trained linear discriminant analysis (LDA) yielded accuracy of 100 %^49^. The cumulative contribution rate of the two components was more than 80 %.

*Task D. (Silent speech of her thought)*. She silently speaks words and phrases yielded by segmentation of sentences which she now wants to talk.

Our RNN-Transformer model was applied to each of single-trial EEGs during silent Japanese answers after the EEG embedding and positional encodings^18^. Note that silent mora ERPs by one student volunteer, but not the patient, were used as the signals in our RNN.

### Data collection and preprocessing

In case of one active electrode (AP-C151-015, Miyuki Giken Co., LTD., Japan), the electrode was affixed to F7 on the scalp according to the International 10-20 System. The average of two earlobe electrodes was used as reference. The EEGs recorded at the electrodes were fed to a wireless amplifier (Polymate Mini AP108, Miyuki Giken Co., LTD., Japan) with 10,000 gain and a notch filter of 60 Hz. The amplified EEGs were sampled at a rate of 500 Hz during an epoch of 2 s preceding and 2 s following each word or phrase presentation. The on-line A/D converted EEG data was immediately transmitted via Bluetooth and stored on a disk in a personal computer.

When using a Muse EEG headband (InterAxon Inc., Toronto, ON, Canada), single-trial EEGs were recorded only at one electrode AF7 with FPz as the reference electrode, being sampled at 256 Hz.

### The network

#### EEG Embedding

This embedding refers to segmentation between 0 and 300 ms then blocking of single-trial EEGs. Each block was blanching to the dot-product layer and the following RNN, as the self-attention in the Transformer. In order to inject some information about the relative or absolute position of the moras in the sequence, also in our model, positional encodings^18^ are added after the EEG embedding.

#### RNN

In order to predict averaged ERPs included in one single-trial EEG, we adopted the recurrent neural network (RNN) whose dynamics is described by:

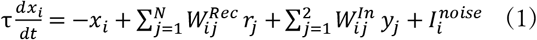

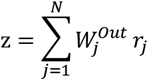

where *r*_*j*_*=tanh(x*_*j*_) represents the activity level of recurrent unit *x*_*i*_ (*i*=1,…,N). The variable *y*_*i*_ represents the input units (*i*=1,2), and *z* is the output. N=800 is the network size, and τ=10 ms is the unit time constant (for more details, see ref.^24^).

The sparse N×N matrix *W*^*Rec*^ represents the recurrent connectivity. The 1 ×N output connectivity vector *W*^*Out*^, the input weight vector *W*^*In*^ and an N×1 random vector *I*^*noise*^ are drawn all from Gaussian distributions. The learning rule for training the plastic recurrent units is based on the RLS (recursive least square)^51^, which was implemented by FORCE (first-order reduced and controlled error) algorithm^52^. The rule was applied to a subset of the recurrent synapses in the network. So, the weight update of the *W*^*Rec*^ is:

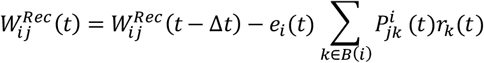

where *B(i)* is the subset of recurrent units presynaptic to unit *i*, and *e*_*i*_ represents the individual error of unit i defined by:

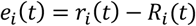

where *r*_*i*_*(t)* is the activity of unit i before the weight update, and *R*_*i*_*(t)* is the “innate” target activity of that unit. *P*^*i*^ is updated by:

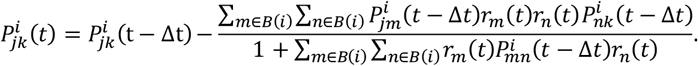

By assigning one single-trial EEG and each of averaged ERPs during silent moras to *st(t)* and *R*_*i*_*(t)*, respectively, and then setting so that

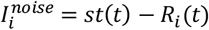

on the basis of the hypothesis on the averaging, we can get approximated solutions of Eq. (1). These solutions were obtained by the Matlab software (ref.^24^, p.926, ‘Supplementary Matlab Routines’).

#### Multi-head attention (MHA)

Our model has MHA structure in the Transformer, where *Q* (query) is assigned to decomposed single-trial EEG, *K* (key) is obtained by the above RNN output for each silent mora ERP, and *V* (value) is given so that the *V* and Affine layer form LSTM (long short-term memory)^29^. The dot-product between the *Q* and the *K* indicates how much the *Q* included each *K*. The relationship among *Q, K* and *V* is given by

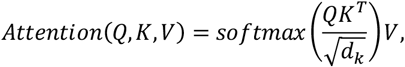

where *d*_*k*_ is the dimension of *Q* and *K*. The number of the heads is equal to that of the blocks. Each block was blanching to the dot-product layer and the above RNN, as the self-attention in the Transformer.

#### Softmax with Loss

Next, “Softmax with Loss” layer calculates Softmax function and cross entropy error. Other loss function can be applied such as MSE Loss and CTC Loss, where the latter Loss can train the neural network to output a sequence of symbols (in this case, moras) given unlabeled time series input^30^. “Mora ID” as teacher signal in Fig. 1c (left) is settled as follows: blanks are assigned to the first and last blocks; dividing the number of the resting blocks by that of moras consisting of word, phrase or (part of) sentence to be silently spoken, moras of the obtained number are given in turn. In our Transformer, MSE Loss yielded the best performance. Lastly, mora probabilities were generated as functions of block number. Tracing mora with the maximum probability at each block yields the decoding of single-trial EEG during silent Japanese speeches.

#### Implementation

##### architecture

The basic architecture of the network, shown in Fig. 1, was modeled after the encoder-decoder neural network, although there are significant modifications.

Each sequence of single-trial EEG data segmented between 0 to 300 ms enters the network through “EEG embedding”. In it, the data are decomposed into temporally overlapping blocks (Fig. 1a). In case of the sampling frequency of 500 Hz, the block size is 200 ms, and the gap between any two adjacent blocks is 2 ms. So, the total number *h* of the blocks is 50 per trial. Given at most 109 silent-mora-related potentials, each ERP and each block minus the ERP are assigned to the signal and noise. Then, eq. (1) is solved, and the solutions, that is, RNN outputs, yield mora ERPs estimated under the noise (Fig. 1b). The RNN was implemented with MATLAB R2022b.

Next, in our Transformer, matrices Q, K and V become each block after positional encoding, the RNN outputs and so that the V and Affine layer construct LSTM, respectively. The LSTM cell is an 800-dimensional vector. The dot-product between the Q and the K indicates how much the Q partly included each K. “Softmax with Loss” (Fig. 1c) calculates Softmax function and minimizes cross entropy error under the teaching signals (mora ID with “blank”). This method was implemented with Python 3.10.9 and PyTorch on an Anaconda Navigator.

## Acknowledgements

We thank the patient and her caregivers for their generously volunteered time and effort, and also Tsukasa Kajihara from Miyuki Giken Kyushu, Co., LTD for supporting the EEG experiments in the patient’s house. We thank Prof. R. Laje and Prof. D. V. Buonomano for the Matlab software of the RNN. This study had been funded by a Grants-in-Aid for Scientific Research on Scientific Research (C) (15K00276) – The Japan Society for the Promotion of Science and TSUMURA&Co. The authors thank several graduates and students for carrying out EEG measurement and EEG data analysis.

## Author contributions

To.Y. and S.-I. K conceived and implemented the decoder and all analyses thereof. S.K. executed the analyses by the decoder. S.T. carried out the preliminary and multi-channel EEG measurement experiment and analyzed the data. S.F. conceived EEG Web interfaces as experimental environment in the future. S.A. and Te.Y. instructed their students concerning EEG measurement experiments and the data analysis. To.Y wrote the manuscript with input from all authors.

## Completing interest

All the authors have no completing interests.

